# Subthreshold electric fields bidirectionally modulate neurotransmitter release through axon polarization

**DOI:** 10.1101/2025.02.22.639625

**Authors:** Aman S. Aberra, Madelyn W. Miles, Michael B. Hoppa

## Abstract

1.

Subthreshold electric fields modulate brain activity and demonstrate potential in several therapeutic applications. Electric fields are known to generate heterogenous membrane polarization within neurons due to their complex morphologies. While the effects of somatic and dendritic polarization in postsynaptic neurons have been characterized, the functional consequences of axonal polarization on neurotransmitter release from the presynapse are unknown. Here, we combined noninvasive optogenetic indicators of voltage, calcium and neurotransmitter release to study the subcellular response within single neurons to subthreshold electric fields. We first captured the detailed spatiotemporal polarization profile produced by uniform electric fields within individual neurons. Small polarization of presynaptic boutons produces rapid and powerful modulation of neurotransmitter release, with the direction – facilitation or inhibition – depending on the direction of polarization. We determined that subthreshold electric fields drive this effect by rapidly altering the number of synaptic vesicles participating in neurotransmission, producing effects which resemble short-term plasticity akin to presynaptic homeostatic plasticity. These results provide key insights into the mechanisms of subthreshold electric fields at the cellular level.

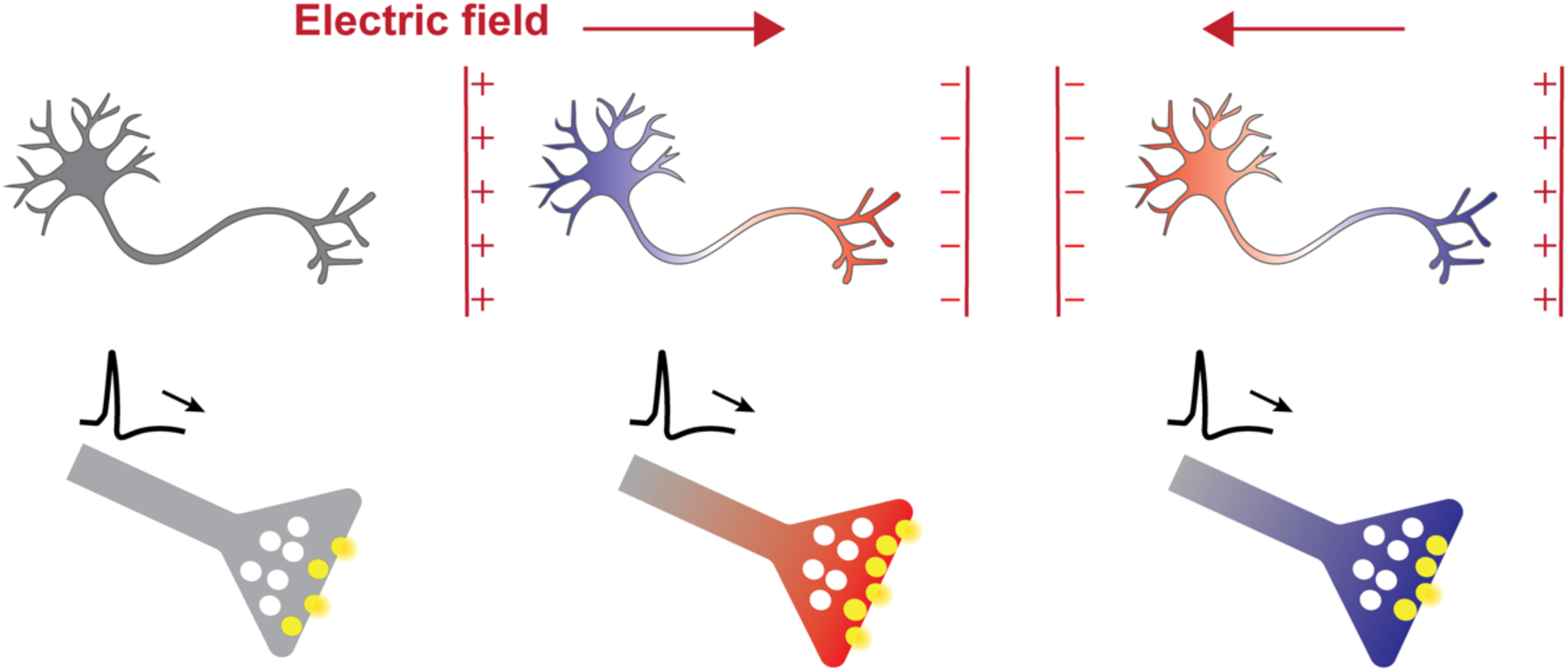

## 2. INTRODUCTION

Extracellular electric fields (E-fields) can be used to modulate neural activity, enabling a myriad of techniques to study functions and treat disorders of the brain. E-fields are generally applied at either suprathreshold strengths, which directly trigger action potentials (APs) in affected neurons, or at subthreshold strengths, which alter excitability and ongoing neural processing. For example, transcranial direct current stimulation (tDCS) is a noninvasive technique in which low-intensity, constant currents are applied via scalp electrodes, producing subthreshold E-fields in the brain. Application of tDCS produces both acute^1^ and lasting changes in brain activity^2^, and tDCS-based therapies have shown promise in treating a variety of maladies including major depressive disorder^3^, fibromyalgia^4,5^, and addiction^6^. However, both the neurophysiological and clinical effects of tDCS often have low efficacy and are highly variable between subjects^2,7–9^. Without a clear understanding of the underlying mechanisms of action, improving efficacy is challenging.

A long history of work in *in vitro* and *in vivo* animal models has probed the cellular and synaptic effects of subthreshold E-fields. Work in acute cortical and hippocampal slice showed subthreshold E-fields polarize neuronal membranes, which can alter spike timing ^10^, evoked and spontaneous post-synaptic potentials ^11–14^, and induction of synaptic plasticity ^15–18^. *In vivo* recordings in freely moving rats found stimulation polarity-dependent changes in spike rates^19^. Detailed modeling of the E-field ^20^, validated by both cadaver^1^ and *in vivo* measurements^21^, found tDCS produces peak E-fields in neocortex of ∼1 V/m with 2 mA applied current. However, significant questions remain when considering mechanism about how these fields act at the subcellular level. Neuronal simulations using one-dimensional cable theory and realistic morphologies predicted that fields of this strength would produce weak somatic membrane polarization (∼0.2 mV in pyramidal cells)^11,22–24^, in agreement with whole-cell patch clamp recordings^25,26^. Interestingly, these models also predict that axons are likely to experience polarization an order of magnitude greater than the soma due to their uniquely elongated morphology and high input impedance. Accordingly, small subthreshold changes in presynaptic resting membrane potential have been shown to alter the AP waveform, Ca^2+^ channel gating, and basal Ca^2+^ concentrations^27–29^. Thus, due to their morphology and function, axons are perfectly positioned to be sensitive to subthreshold E-fields. Unfortunately, direct measurements of E-field-induced axonal polarization and its resulting impact on synaptic transmission are not possible using traditional electrophysiology due to the technique’s low spatial resolution, its need to disrupt the cell membrane and cytosol with a micropipette, and its susceptibility to artifacts caused by changes in surrounding E-fields.

To overcome these limitations, we deployed genetically encoded optical indicators of voltage, glutamate, and Ca^2+^ to probe the effects of uniform, subthreshold E-fields on synaptic transmission in cultured hippocampal neurons. We found subthreshold E-fields produced rapid and dramatic facilitation or inhibition of glutamate release depending on the E-field intensity and polarity. Moreover, we provide evidence that this rapid facilitation is generated by axonal depolarization and a basal Ca^2+^-dependent increase in the number of synaptic vesicles in the readily releasable pool (RRP), rather than changes in AP-evoked Ca^2+^ influx and vesicular release probability. Our results shed light on the fundamental biophysical mechanisms of subthreshold E-field stimulation and support a role for modulation of synaptic transmission in the physiological effects of tDCS.

## 3. RESULTS

### 3.1. Subcellular distribution of membrane polarization by E-field stimulation

To study the axonal and synaptic mechanisms of subthreshold electric field (E-field) stimulation, we applied spatially uniform E-fields to cultured rat hippocampal neurons sparsely labeled with genetically encoded voltage indicators (GEVIs) (**Figure 1A**). Uniform E-fields were generated using a custom stimulation chamber consisting of two pairs of parallel platinum-iridium electrodes lowered onto glass coverslips with cultured neurons (**Figure S1**). We used pharmacological blockers of excitatory synaptic transmission (CNQX/APV) to silence network activity, minimizing background fluctuations in membrane potential and enabling us to isolate the direct subcellular response to applied E-fields.

**Figure 1.**
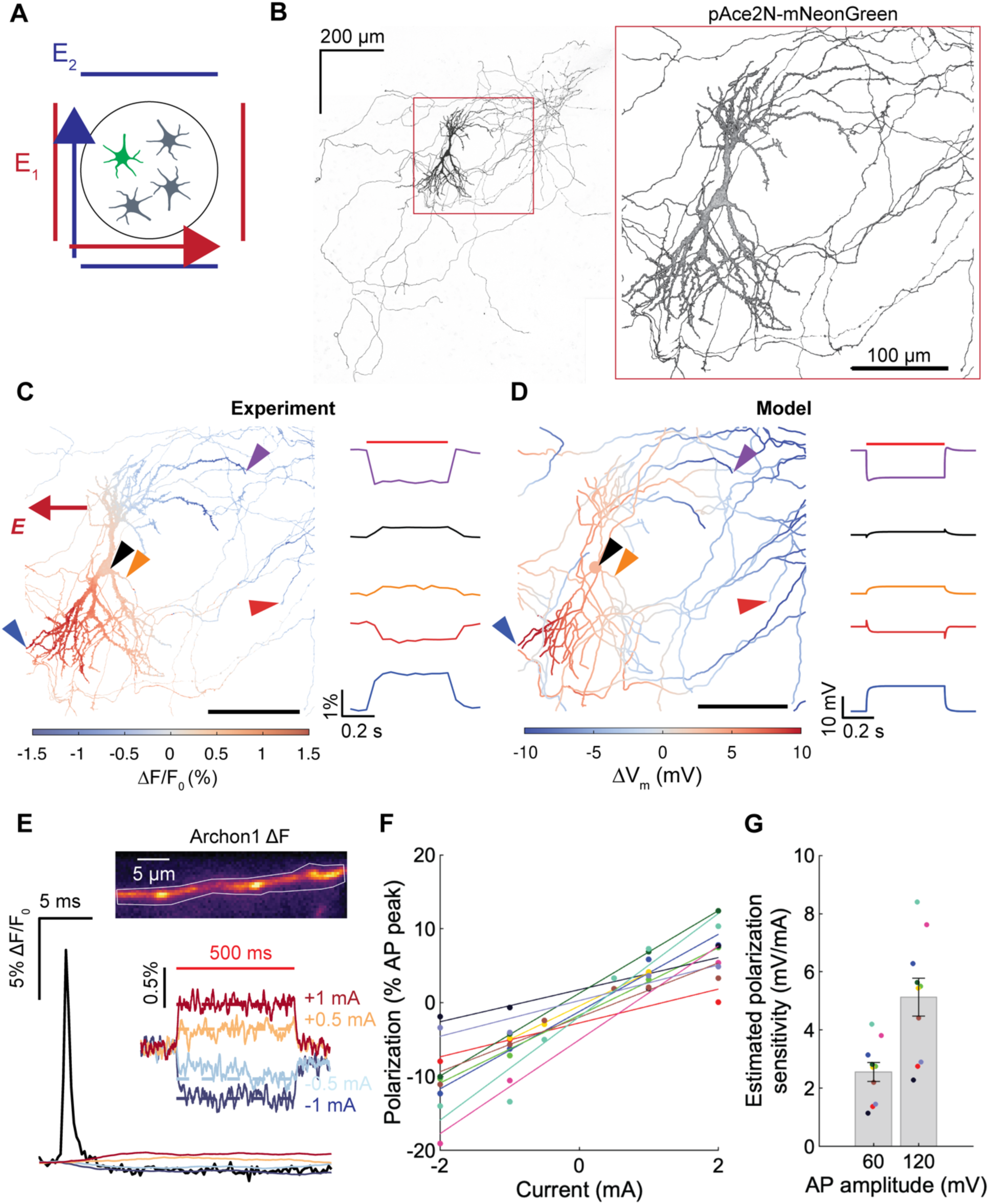
Optical measurement and modeling of E-field-induced membrane polarization. **A)** Schematic of stimulation electrodes enabling 2D control of uniform E-field direction applied to hippocampal cultured neurons sparsely transfected with GEVI pAce2N-mNeon. **B)** Max projection of tiled confocal z-stack of single neuron expressing pAce2N-mNeon (left) and zoom in of field of view (FOV) used for polarization imaging with widefield microscope (right). **C)** Measured steady**-**state change in fluorescence for 500 ms subthreshold E-field (2 mA, ∼110 V/m), with direction indicated by arrow (left), and fluorescence traces from example regions of interest (ROIs) (right). ROIs indicated by colored arrows with soma in black, dendritic ROIs in purple/blue, and axonal ROIs in orange/red. Images collected at 20 Hz on CMOS camera. **D)** Simulated steady-state change in membrane voltage (left) and voltage traces from same ROIs as B (right). **E)** High-speed (2 kHz) imaging of axonal AP waveform with Archon1. Example AP waveform overlaid with response to 500 ms, subthreshold uniform E-field stimulation at varying intensities (bottom). Subthreshold polarization responses filtered using 20 ms wide quadratic Savitsky-Golay filter. Top inset shows mean fluorescence change (Δ*F*) during AP recording trial. Right inset shows zoom in of polarization responses. **F)** Steady-state polarization relative to AP amplitude at each subthreshold stimulus intensity (circles) with linear fit overlaid (solid lines). **G)** Estimated polarization sensitivity (mV/mA) (slope of lines in F) assuming AP peak is either 60 mV or 120 mV (*n* = 10 cells).

We first measured the spatial distribution of changes in membrane potential (polarization) induced by 500 ms E-field pulses using pAce2N-mNeonGreen^30,31^, a positively tuned fluorescence resonance energy transfer (FRET)-opsin genetically encoded voltage indicator (GEVI), on a widefield fluorescent microscope. Functional imaging data (20 Hz) were then co-registered with high resolution structural imaging of the same neuron on a spinning disk confocal microscope (**Figure 1B**). The uniform E-field generated a non-uniform membrane polarization distribution, with depolarization at the cathodal (negative) and hyperpolarization at the anodal (positive) end of the E-field (**Figure 1C**). Additionally, we reconstructed the neuron’s morphology and simulated the response to a uniform E-field of the expected orientation, finding broad agreement between the experiment and model (**Figure 1D**), even when rotating the E-field (**Figure S2**). As expected from cable theory, steady-state polarization within a compartment was a function of both the compartment’s local coupling to the E-field, related to its orientation and cable properties, and axial current redistribution between compartments^23,32^.

While GEVIs have limited ability to measure absolute voltage changes, we calibrated the polarization response to the subthreshold E-field by comparing induced changes in voltage with action potential (AP) waveforms recorded at 2 kHz. Here, we used the GEVI Archon1 due to its rapid kinetics and linearity across the physiological voltage range^33^. **Figure 1E** shows an example axonal recording with the polarization response to varying E-field intensities overlaid with the AP waveform. We measured AP waveforms and polarization responses from one <40 µm axonal region per neuron. Steady-state polarization across the range of E-field intensities used (<2 mA) was approximately linear (*R*^2^ > 0.85 for all axons, linear fit), with sensitivities of 4.4 ± 0.5% of the maximal AP amplitude per mA (n = 10 axons; all statistics are mean ± SEM unless specified otherwise) (**Figure 1F**). Using lower and upper bound estimates of axonal AP amplitudes ^34,35^, we estimated polarization sensitivities of 2.6 ± 0.3 to 5.1 ± 0.6 mV per mA, respectively (**Figure 1G**), with polarization time constants of 9.2 ± 1.1 ms (**Figure S3**). The polarization sensitivity was not correlated with AP amplitude indicating the variations in responses between axons were not related to reporter expression or trafficking and likely reflected differential E-field sensitivity (**Figure S4**).

Thus, we established a platform for measuring subcellular responses within single neurons to E-fields; voltage imaging indicated polarization distributions and time courses largely agree with theoretical predictions for subthreshold uniform E-fields based on cable theory. Next, we investigated the impact of these subthreshold voltage changes on glutamatergic neurotransmitter release.

### 3.2. Subthreshold DC, E-field stimulation acutely facilitates or inhibits evoked glutamate release depending on stimulus polarity and intensity

We next asked what impact these subthreshold axonal voltage changes have on neurotransmission. We labeled presynaptic boutons using synapsin-mRuby and quantified neurotransmitter release using a highly sensitive glutamate sensor, iGluSnFR3^36,37^ (**Figure 2A**). To evoke action potentials, we used a bipolar stimulation electrode positioned near the soma of individual iGluSnFR3-expressing neurons, again while pharmacologically silencing network activity. We measured AP-evoked neurotransmitter release in single boutons before, during, and after application of a 10 sec, subthreshold uniform E-field, using the same setup seen in Figure 1. We found rapid and dramatic facilitation or inhibition of neurotransmitter release during application of the subthreshold E-field, depending on the E-field polarity (**Figure 2B–C**). We characterized the dose–response relationship by varying the applied current intensity (±0.1, 0.5, and 1 mA). Subthreshold, DC E-fields bidirectionally impacted the magnitude of glutamate release at all intensities (**Figure 2D**). The peak of the stimulus averaged iGluSnFR3 response (Δ*F*/*F*_0_) increased 86.8 ± 5.0% at 0.5 mA and decreased 22.8 ± 1.6% at −0.5 mA (mean ± SEM; **Figure 2D**). Peak glutamate release was also significantly modulated by intensities as low as 0.1 mA, increasing by 8.8 ± 1.3% and decreasing by 7.2 ± 1.1% (**Figure 2D**). Taking advantage of the high signal to noise ratio of iGluSnFR3, we analyzed responses within bouton at each intensity to identify significant responders at each intensity. Given the stochastic nature of synaptic release and highly variable release probability between boutons^38^, we controlled for multiple comparisons and false discoveries using a conservative 1% cutoff (*see Methods*). We found 0.1 mA DC stimulation modulated 5–8% of synapses, while two thirds of synapses were facilitated and one third were inhibited at ±0.5 and ±1 mA (**Figure 2E**). Interestingly, we noticed a subset of synapsin-mRuby+ synapses that failed to release neurotransmitter before E-field stimulation on any trial but noticeably became active or “unsilenced” during subthreshold E-field stimulation and vice versa (**Figure 2F**). Thus, presynaptic boutons are highly sensitive to subthreshold E-field stimulation, which rapidly and bidirectionally modulates neurotransmitter release.

**Figure 2.**
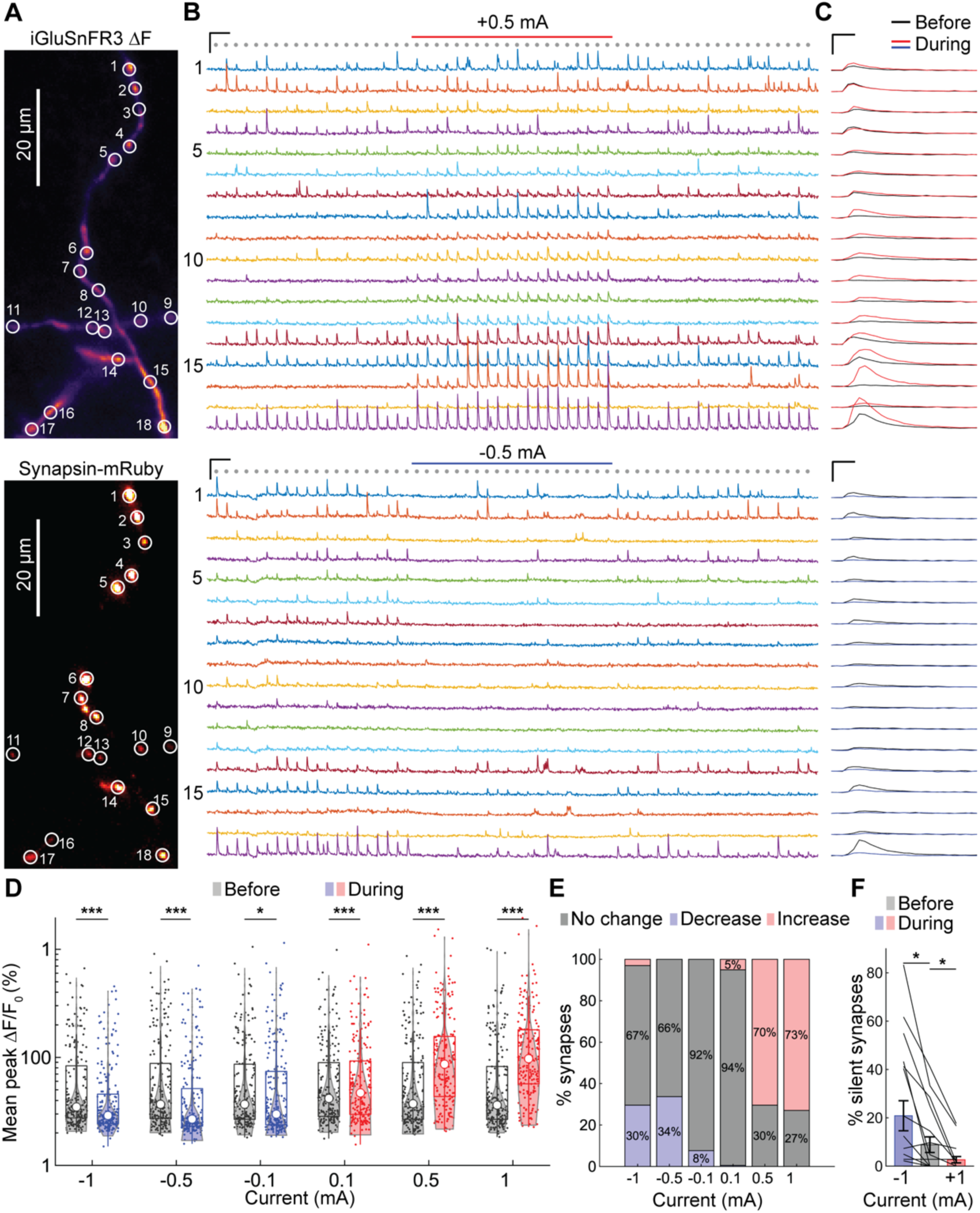
Subthreshold DC, E-field stimulation acutely facilitates or inhibits evoked glutamate release depending on stimulus polarity and intensity. **A)** Example cell expressing iGluSnFR3 (top) and synapsin-mRuby (bottom) with ROI outlines overlaid. **B)** Fluorescence (Δ*F*/*F*_0_) traces from ROIs shown in A during 2 Hz AP train evoked by bipolar electrodes (indicated by grey circles) with 10 sec DC, uniform E-field applied after 10.25 sec (250 ms before the next AP, indicated by red or blue bar above fluorescence traces). Facilitating polarity (+0.5 mA) shown on top, inhibiting (−0.5 mA) shown on bottom. Scale bars = 1 sec, 100% Δ*F*/*F*_0_. **C)** Stimulus averaged AP-evoked responses of 20 APs before and during DC E-field stimulation in each ROI overlaid. Scale bars = 40 ms, 100% Δ*F*/*F*_0_. **D)** Distribution of peaks within period (before or during stimulation at each intensity) shown as violin plot, with overlaid box and whisker plot. Individual points are mean of 40 APs (20 APs, 2 trials) within period for each ROI (*n* = 196 boutons, 8 cells). * *p* < 0.05, ** *p* < 0.01, *** *p* < 0.001, post-hoc Tukey-Kramer on distributions during (red/blue) compared to before DC stimulation after two-way RM-ANOVA. Neurons in culture have no intrinsic orientation preference and thus predictable polarization, so we used the responses to the strongest subthreshold E-field (+1 and −1 mA) to designate the facilitating E-field direction “positive” for each bouton and pool all single bouton data (see Methods). **E)** Fraction of significantly modulated synapses during DC E-field at each intensity, determined using Wilcoxon signed-rank tests within synapse corrected for multiple comparisons using the Benjamini and Hochberg procedure with false discovery rate set to 1%. **F)** Change in proportion of silent synapses before and during +1 mA (facilitating) DC. Silent synapses defined as synapses that failed to produce responses over 1 STD of baseline, i.e. release probability below our detection limit (<1/40 APs = 1.25%). Data are represented as mean ± SEM (*n* = 18 cells, 534 boutons, * *p* = 0.016, Wilcoxon signed-rank).

### 3.3. Subthreshold E-fields facilitate release by increasing the size of the readily releasable pool of synaptic vesicles

According to the quantal model of synaptic release, the strength of a synapse is proportional to the product of the number of release sites, *N*, also known as the readily releasable pool of vesicles (RRP), and their probability of release *P*_*v*_^39,40^. Therefore, we wanted to determine which of these two variables the subthreshold E-field acted upon to rapidly change presynaptic strength. To distinguish between changes to RRP or *P*_*v*_, we measured paired pulse ratio (PPR) of iGluSnFR3 responses at baseline and in the presence of 2 s duration (+50 V/m) E-field stimulation (**Figure 3A**). If E-field polarization facilitated release by increasing *P*_*v*_ without increasing *N*, we would expect the PPR to be reduced, since fewer release-ready vesicles would remain to fuse for the second AP^41^. However, if *N* were increased, then PPR could remain the same, as seen in some types of presynaptic homeostatic plasticity^42^. We found that under control conditions, our boutons exhibited a facilitating PPR of 1.26 ± 0.08 (**Figure 3B–D**). Application of the subthreshold E-field augmented the magnitude of release on the first action potential by ∼90% but also maintained a nearly identical PPR of 1.25 ± 0.06 (n = 7 cells, 152 boutons) (**Figure 3B–D**). These data strongly support the model in which the subthreshold E-field acts on the size of the RRP, rather than *P*_*v*_, to modulate the magnitude of neurotransmitter release.

**Figure 3.**
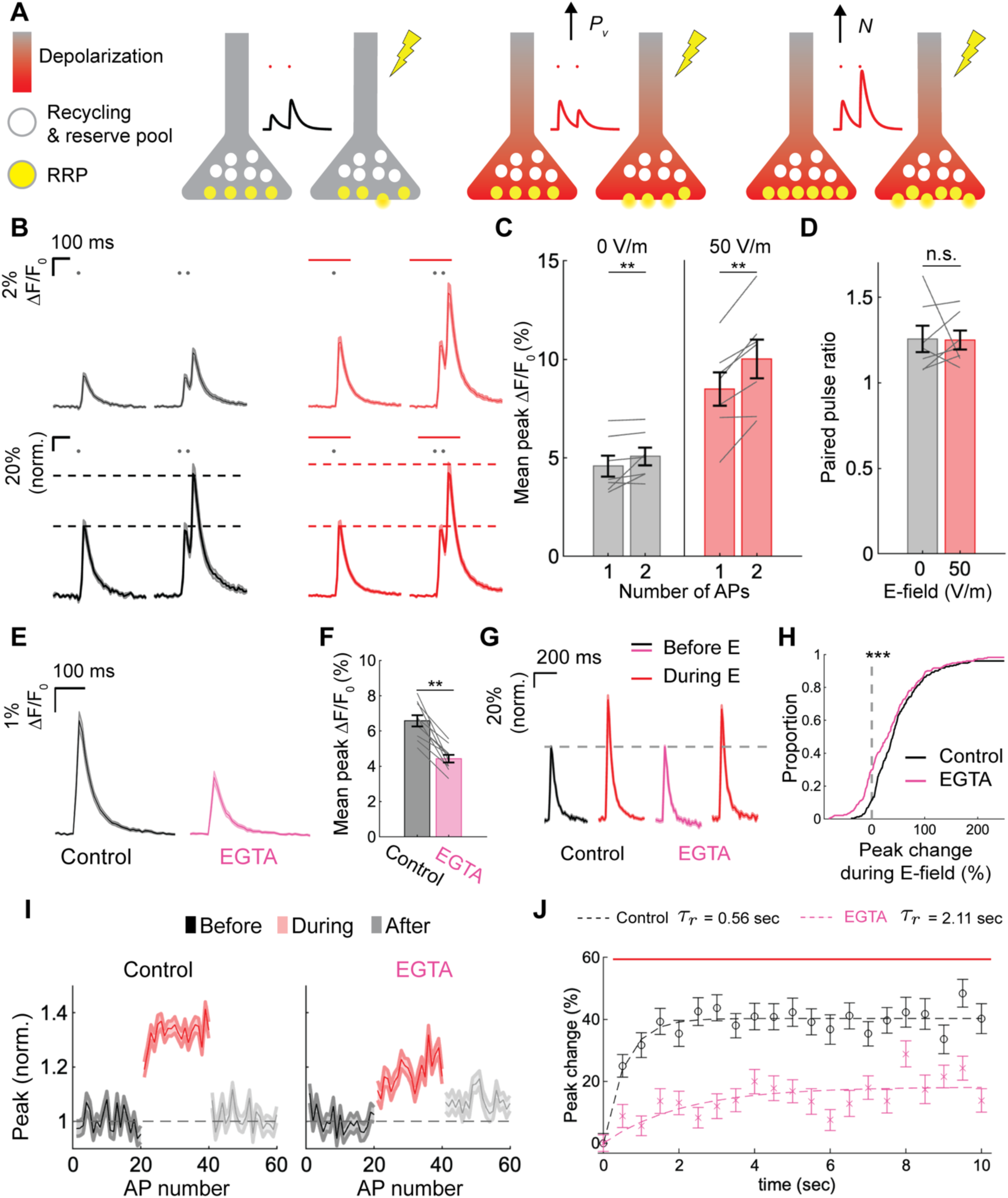
Facilitation by subthreshold, DC E-field is mediated by a basal Ca^2+^-dependent increase in the readily releasable pool. **A)** Cartoon demonstrating two possible mechanisms for E-field-induced facilitation and predicted response to paired pulse stimulation. Under baseline conditions (left pair of presynaptic terminals), an AP (lightning bolt) triggers vesicle release with probability proportional to the vesicular release probability (*P*_*v*_) and number of vesicles in the readily releasable pool (RRP). Depolarization by a subthreshold E-field could increase *P*_*v*_, making vesicles already in the RRP more likely to fuse upon arrival of an AP and increasing net neurotransmitter output (middle pair of terminals). Alternatively, depolarization could recruit more vesicles to the RRP, leading to increased neurotransmitter output without requiring an increase in *P*_*v*_ (right pair of terminals). **B)** Mean iGluSnFR3 fluorescence traces (Δ*F*/*F*_0_) in response to single or paired pulse (20 Hz) stimulation in absence (black, left) or presence (red, right) of 50 V/m, 2.1 sec E-field pulse (indicated by red bar). Traces in top row are unnormalized, while traces in bottom row are normalized to the single pulse trials within synapse and E-field condition (mean ± SEM, n = 7 cells, 152 boutons). Dashed lines indicate peaks used for quantifying paired pulse ratio. **C)** Effect of 50 V/m E-field on mean peak evoked iGluSnFR3 (Δ*F*/*F*_0_) for single and paired pulse stimuli (**, p < 0.01, post-hoc Tukey-Kramer after two-way RM ANOVA). Note: peak of paired pulse stimulus corrected for summation with decay of first stimulus by subtracting fluorescent ((Δ*F*/*F*_0_) value preceding second stimulus in 20 Hz paired pulse. **D)** Effect of 50 V/m E-field on paired pulse ratio, calculated with peak Δ*F*/*F*_0_ response to 2 AP divided by peak Δ*F*/*F*_0_ response to 1 AP (p = 0.71, Wilcoxon signed-rank) within bouton before averaging across boutons within each cell. **E)** Effect of incubation in 100 µM EGTA-AM on mean AP-evoked iGluSnFR3 response Δ*F*/*F*_0_(mean ± SEM, n = 10 cells, 233 boutons). **F)** Mean peak iGluSnFR3 Δ*F*/*F*_0_ response in control and EGTA (**, p = 0.002, Wilcoxon signed-rank). **G)** Mean AP-evoked iGluSnFR3 response (Δ*F*/*F*_0_) normalized to peak before E-field in control and after incubation in EGTA. **H)** Cumulative distribution plots of percent change in glutamate release (peak Δ*F*/*F*_0_) during E-field for facilitating subthreshold E-field (±1 mA, ∼50 V/m) in control (black) and after EGTA (pink). (*** p = 0.001, Wilcoxon signed rank). **I)** Peak iGluSnFR3 Δ*F*/*F*_0_ for each AP within 2 Hz train before (1–20, black), during (21–40, red), and after (41–60, gray) subthreshold E-field stimulation, normalized to mean before E-field within bouton, in control and after EGTA. **J)** Percent change in peak Δ*F*/*F*_0_ for facilitating subthreshold E-field relative to last AP before E-field for control (black) and EGTA (magenta), with overlaid monoexponential fit (dashed lines).

One of the key drivers to modulating the size of the RRP are small changes to basal Ca^2+^ levels in the presynapse^43–45^. To test this hypothesis, we first used a slow, high-affinity Ca^2+^ chelator, ethylene glycol tetraacetic acid using its acetoxymethyl ester form for intracellular delivery (EGTA-AM). EGTA restricts Ca^2+^ diffusion to microdomains near sources (e.g., voltage gated Ca^2+^ channels during APs), reducing and slowing changes in basal Ca^2+^ in the cytosol. Using iGluSnFR3 to measure AP-evoked glutamate release with a 2 Hz AP train, EGTA-AM (100 µM) reduced baseline glutamate release 31.85 ± 0.72% across synapses (**Figure 3E–F**). After loading with EGTA, mean facilitation during a 10 sec subthreshold E-field (same protocol as in **Figure 2**) was reduced by 31.0 ± 10.7% relative to control (**Figure 3G–H**). In our previous analysis (**Figure 2**), we averaged the peaks of all AP responses during the 10 sec E-field; however, when we analyzed the mean peak response to each AP in the train relative to the pre-E-field average within synapse, we observed facilitation during the E-field increased with each AP before reaching a steady state within ∼3 sec. Interestingly, this buildup of facilitation was also slowed by EGTA (**Figure 3I–J**). The time course of peak modulation over time was captured well by a monoexponential function with time constants of 0.56 sec (*R*^2^ = 0.89) in control and 2.11 sec (*R*^2^ = 0.49) in EGTA (**Figure 3J**). The time course and EGTA sensitivity of facilitation is consistent with facilitation by the subthreshold E-field being mediated by small changes in basal Ca^2+^ that reach a steady-state during application of the E-field, which would go undetected by genetically encoded Ca^2+^ indicators due to slow kinetics and likely tens of nM changes. Taken together, these data support a model in which the subthreshold E-field enhances RRP size through increases in basal Ca^2+^.

### 3.4. Facilitation by subthreshold, DC E-field is not explained by AP broadening or enhanced AP-evoked Ca influx

While subthreshold E-field modulation of RRP size surprisingly seemed to account for the changes in glutamate release we observed, we also wanted to test the role of changes in *P*_*v*_ that might occur by modulation of ionic currents shaping the AP waveform. Inactivating axonal voltage-gated K^+^ (Kv) channels, specifically D-type Kv1 channels, has been associated with depolarization-induced analog facilitation of synaptic transmission^46^. To determine if this mechanism could explain the E-field induced facilitation or inhibition, we first measured axonal AP waveforms before and during application of a 10 sec, DC E-field with both polarities (±1 mA), mimicking the protocol used in our glutamate imaging experiments. Surprisingly, we found that neither AP width nor the waveform in general were affected by either E-field polarity (**Figure 4A–B**). This is likely due to the modest changes to membrane voltage elicited by the weak E-fields. In contrast, even partial inhibition of Kv1 channels with 50 nM alpha-dendrotoxin (DTX) robustly broadened AP width by 33.6 ± 6.1% (**Figure 4B**), demonstrating our ability to detect AP broadening by Kv1 blockade. Previously, we found that activity-dependent inactivation of Kv1 channels in hippocampal axons requires the Kvβ1 subunit ^47^. Knocking down K_v_β1 expression also failed to prevent facilitation using an E-field (**Figure S5**). Taken together, these data suggest that the strong facilitation of glutamate release by subthreshold E-field stimulation is not mediated by changes in axonal AP waveform shape, including inactivation of low-threshold Kv1 channels.

**Figure 4.**
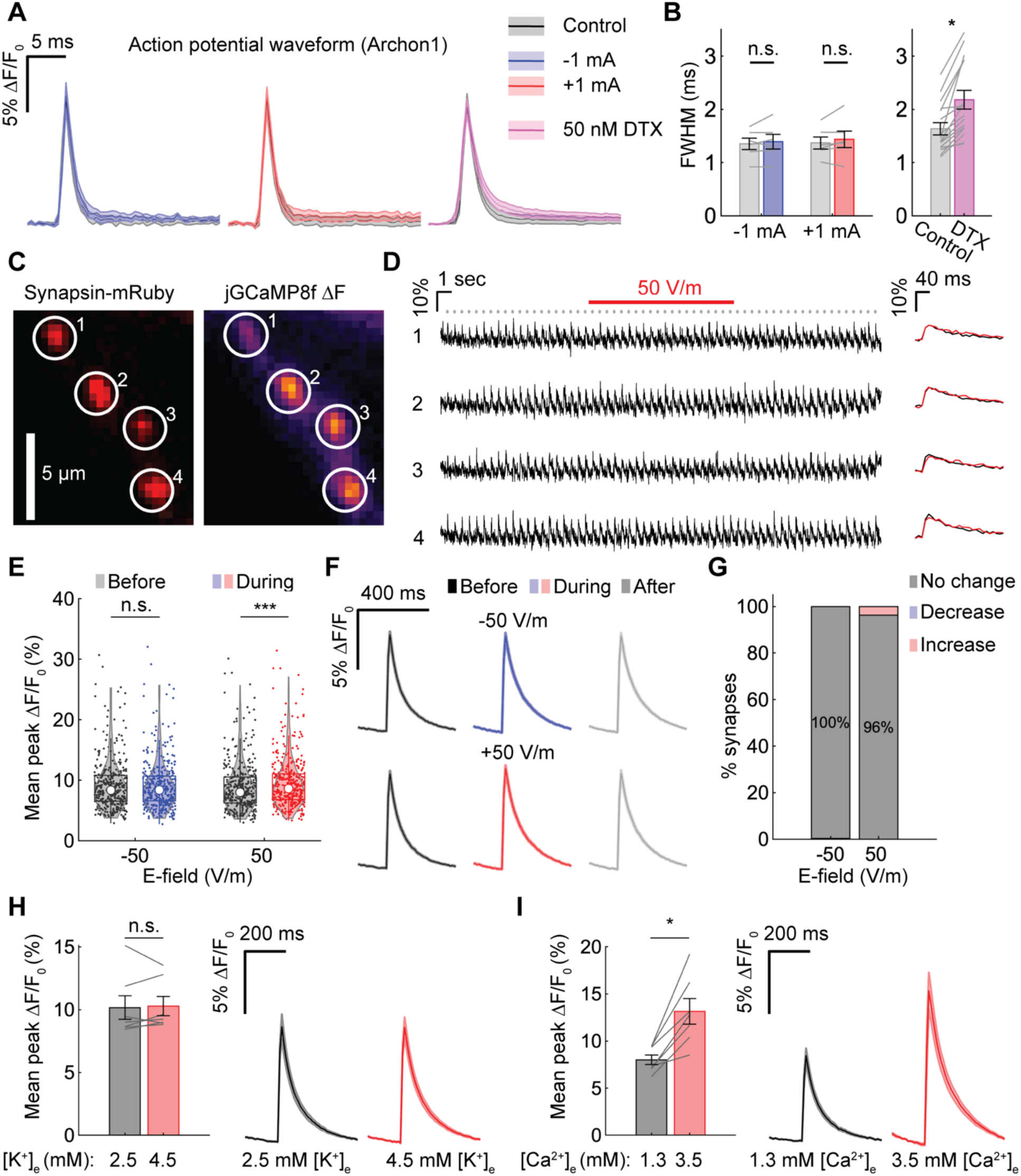
Facilitation by subthreshold, DC E-field is not explained by AP broadening or enhanced AP-evoked Ca influx. **A)** Fluorescence traces of axonal AP waveforms (mean ± SEM) measured before (control) or during 10 sec hyperpolarizing (−1 mA, left) or depolarizing (+1 mA, middle) DC, E-field, or before and after application of 50 nM DTX (right). AP waveforms within axon averaged from 20 APs (±1 mA) or 50 APs (50 nM DTX) evoked in 2 Hz train. **B)** Full-width half-maximum of AP waveforms before and during ±1 mA stimulation (left) (*n* = 6 cells, *p* > 0.05 for all main effects and interactions of two-way RM ANOVA) and before and after application of 50 nM DTX (right, *n* = 16 cells, * *p* < 0.001, Wilcoxon signed-rank). **C)** Example fluorescence images of representative axon branch showing synapsin-mRuby (left, baseline fluorescence) and jGCaMP8f (right, mean Δ*F* of 60 APs evoked at 2 Hz). **D)** Fluorescence traces from four boutons in C) during 2 Hz AP train evoked with E-field stimulation via perpendicular pair of electrodes (1 ms pulses, indicated by grey circles). Subthreshold, DC uniform E-field applied for 10 sec after 10.25 sec (250 ms before the next AP, indicated by red bar). **E)** Distribution of peaks within period (before or during stimulation at each intensity) shown as violin plot, with overlaid box and whisker plot. Individual points are mean of 40 APs within period for each ROI (*n* = 366 boutons, 14 cells). *** *p* < 0.001, post-hoc Tukey-Kramer on distributions during (red/blue) DC compared to before DC after two-way RM-ANOVA. **F)** Fraction of significantly modulated synapses during DC E-field at each intensity, determined using the Benjamini and Hochberg procedure with false discovery rate set to 1%. **G)** Fluorescence traces of AP-evoked Ca influx (mean ± SEM) averaged before, during, and after stimulation with dc, E-field (−50 or +50 V/m). **H)** Effect of increasing [K^+^]_e_ on evoked jGCaMP8f peak (left) and fluorescence trace (Δ*F*/*F*_0_, right) (n = 7 cells, 208 boutons, *p* = 0.58, Wilcoxon signed-rank). Stimulation protocol consisted of 40 APs delivered at 2 Hz. **I)** Effect of increasing [Ca^2+^]_e_ on evoked jGCaMP8f peak (left) and fluorescence trace (Δ*F*/*F*_0_, right) (*n* = 7 cells, 172 boutons, * *p* = 0.016, Wilcoxon signed-rank). Same stimulation protocol as H). Note: independent samples for statistical tests in H–I are cells, using responses averaged across boutons, rather than individual boutons.

Subthreshold polarization could also alter *P*_*v*_ by acting directly on voltage gated Ca^2+^ channels and enhancing or suppressing AP-evoked Ca^2+^ influx. Indeed, given the enhancement of neurotransmitter release by a facilitating E-field (+118%), we would predict more than a 30% increase in Ca^2+^ influx if this were the primary mechanism driving increased synaptic strength ^48^. We used jGCaMP8f to measure the effect of subthreshold E-field stimulation on AP-evoked cytosolic Ca^2+^ influx within synapses marked with synapsin-mRuby, using the same stimulation protocol and analysis approach used previously with iGluSnFR3 (**Figure 4C–D**). AP-evoked jGCaMP8f signals were similar before, during, and after application of the 10 sec, 50 V/m E-field. Across all cells, peak jGCaMP8f (Δ*F*/*F*_0_) slightly increased (7.7 ± 0.4%) for the assigned facilitatory E-field direction (+50 V/m, +1 mA) for a small minority of boutons and was not significantly altered by the opposite direction (**Figure 4E–F**). Across all boutons, statistically significant responders comprised only 4% of synapses (14/366 synapses, 14 cells) (**Figure 4G**).

This suggested that small depolarization facilitate, whereas hyperpolarizations depress release, without altering AP-evoked Ca^2+^ influx. We wanted to confirm this finding independent of E-field stimulation, so we tested how shifting the resting membrane potential affects glutamate release by modifying the extracellular K^+^ concentration ([K^+^]_e_). Increasing [K^+^]_e_ from 2 to 4.5 mM (∼5 mV depolarization estimated using the Nernst equation) facilitated glutamate release, increasing peak iGluSnFR3 responses by 24.9 ± 3.4% within bouton, while lowering [K^+^]_e_ to 1 mM (∼5 mV hyperpolarization) inhibited release, reducing peaks by 33.5 ± 1.4% (**Figure S6**). Thus, these results supported our hypothesis that depolarization of resting membrane potential facilitated neurotransmission. Like with subthreshold E-field stimulation, we found slightly increasing [K^+^]_e_ also produced negligible effects on AP-evoked Ca^2+^ influx (**Figure 4H**), despite causing a 25% increase in glutamate release (**Figure S6C–D**). To verify we could detect relevant increases in axonal Ca^2+^ influx, we tested the effect of raising extracellular Ca^2+^ ([Ca^2+^]_e_), a standard method of increasing driving force and Ca^2+^ influx. Indeed, raising [Ca^2+^]_e_ from 1.3 to 3.5 mM enhanced peak jGCaMP8f by 63.2 ± 2.2% (**Figure 4I**). In sum, these results indicate subthreshold depolarization by the E-fields we applied facilitated neurotransmission without altering the presynaptic AP waveform or AP-evoked Ca^2+^ influx. Taken together, these results support a model where subthreshold E-Fields alter synaptic strength through changes in the RRP rather than *P*_*v*_.

## 4. DISCUSSION

Most therapeutically-applied E-fields appear spatially uniform at the level of single neurons^49^, particularly as the distance from source to target increases. However, the unique morphology of neurons, especially axons, results in highly non-uniform polarization. Given the many non-linearities that exist between, voltage, Ca^2+^ and vesicle fusion in presynaptic compartments ^34,50,51^, axons are well positioned to be sensitive to modulation by subthreshold E-fields.

We took advantage of several optical indicators that enabled us to non-invasively measure membrane polarization and quantify presynaptic function during subthreshold, DC E-field stimulation. First, our voltage imaging revealed the spatial distribution of membrane polarization in an electric fields: the uniform E-field depolarized compartments near the cathodal (negative) end and hyperpolarized compartments near the anodal (positive) end, in agreement with cable models^11,23^. These data were consistent with mesoscopic voltage sensitive dye imaging in acute brain slices^25^, while for the first time, providing subcellular resolution neuronal polarization. Additionally, we found that subthreshold, DC E-fields can bidirectionally modulate glutamate release via presynaptic membrane polarization depending on the intensity and polarity of local voltage changes. Subthreshold depolarization in central synapses generally enhances synaptic transmission^52–56^, while hyperpolarization suppresses transmission, with some exceptions^57,58^. By monitoring glutamate release simultaneously from several synaptic boutons, we found depolarizing E-fields facilitate and hyperpolarizing E-fields inhibit synaptic vesicle fusion. We provide evidence that this facilitation may be mediated by rapid (∼100 ms scale) recruitment of synaptic vesicles to the RRP, as opposed to changes in P_v_ that can occur by changes to the AP waveform shape or Ca^2+^ channel gating. Mechanistically, increased basal Ca^2+^ may enhance the RRP by accelerating the rate of synaptic vesicle docking, in line with recent reports of rapid activity- and Ca^2+^-dependent vesicle docking ^45^. Thus, we have identified a new mechanism by which subthreshold E-fields can strongly control synaptic transmission.

We found that subthreshold E-fields were able to consistently and bidirectionally modulate synapses independent of their baseline strength, which again is more easily explained if changes occur at the level of the RRP. An advantage of this form of synaptic modulation is that it alters synaptic strength while maintaining the short-term plasticity (STP) critical for circuit function^59^. Intriguingly, our findings closely parallel an essential form of synaptic plasticity known as presynaptic homeostatic plasticity (PHP). PHP is an evolutionarily conserved form of homeostatic control that is expressed in invertebrate^60^ and mammalian synapses^61,62^. PHP can be induced within minutes and dramatically increase synaptic vesicle fusion by up to 200%^60,63^. This plasticity modulates neurotransmitter release depending on postsynaptic sensitivity to neurotransmitter, maintaining synaptic communication, while also preserving STP dynamics. Although not all mechanisms controlling or initiating PHP are known, two conserved mechanisms for PHP have been identified. The first is a Na^+^ leak current generated by epithelial sodium channels (ENaCs), which elevates resting membrane potential^64^. The second involves proteins essential to vesicle docking and priming to form the RRP^63^. Thus, there are uncanny parallels between synaptic modulation by subthreshold E-fields and PHP: both are initiated by “analog” changes in resting membrane potential, and both change synaptic output by modulating the RRP.

One of the most powerful and therapeutically-relevant aspects of noninvasive brain stimulation techniques such as tDCS is their ability to induce lasting changes in cortical excitability after stimulation has ended, suggesting they recruit synaptic plasticity mechanisms ^2^. Many research groups have investigated links between tDCS and Hebbian forms of plasticity ^18^; based on the mechanism and time course of the synaptic changes we observed, we suggest that subthreshold E-fields are able to initiate homeostatic plasticity mechanisms, allowing these fields to exert lasting effects on brain circuits.. This is intriguing as impairments in homeostatic plasticity are potentially causative in a number of neurological diseases and are a currently underexplored therapeutic target ^65^. Taken together this suggests that, depending on the strength and duration of the E-field, subthreshold electrical stimulation could engage multiple forms of synaptic plasticity.

We studied the effects of subthreshold E-fields in cultured hippocampal neurons, providing ideal conditions for imaging from single synaptic boutons. We found glutamate release was modulated in 5–8% of synapses at the lowest E-field intensity used (∼5.5 V/m), yielding average sensitivity of glutamate release of 1.3–1.6% per V/m, assuming the effects can be linearly extrapolated. However, due to differences in morphology, myelination, and ion channel distribution that likely impact overall E-field sensitivity^22^ (i.e., mV polarization per V/m E-field), it is challenging to directly infer effects on human neurons from our experiments in rodent hippocampal culture. At the same time, dosing based on a specific physiological effect—in this case, induced axonal polarization—rather than the absolute E-field amplitude may be more relevant when comparing doses across preparations or delivering effective therapeutic doses across subjects. Indeed, suprathreshold stimulation techniques like transcranial magnetic stimulation and deep brain stimulation are already individualized by adjusting the stimulation intensity relative to a target physiological or behavioral response (e.g. % of motor threshold), an adjustment that is necessary due to variability in individual anatomy and brain state.

Our results also raise several important questions. How does the presynaptic modulation we observed interact with postsynaptic polarization to broadly affect synaptic transmission? How specific are these effects on information processing by glutamatergic synapses at the circuit or regional level, given the distribution of local and afferent axonal orientations? Are inhibitory or neuromodulatory axons similarly modulated by subthreshold E-fields? These questions may be readily addressed using optical approaches to measure axonal physiology and structure, as demonstrated in this study.

Here, we report the first detailed subcellular measurements of the neuronal response to subthreshold E-fields using optical physiology. We found that subthreshold changes in voltage are sufficient to bidirectionally modulate synaptic strength while maintaining essential aspects of short-term plasticity. Our findings could inform development of therapies for psychiatric and neurological disorders using long duration subthreshold E-fields, providing novel avenues for treatment.

## ACKNOWLEDGEMENTS

We thank Dr. Lauren Panzera and Dr. Michelle Gleason for help in preparing neuronal cultures and conducting imaging experiments and Zherong Zhang for help with neuron morphology reconstruction. We also thank the rest of the Hoppa Lab for helpful discussions and feedback. We also acknowledge Emily Dueñas Cornell Du Houx for help with visualizations, as well as Dr. Allan Gulledge, Dr. Robert Hill and Hill lab members, and Dr. Angel Peterchev for feedback on early versions of this work. We thank Ann Lavanway for her expertise and support in the use of the spinning disk confocal microscope. This work was supported by a Neukom Fellowship and CompX grant from the Neukom Institute for Computational Science at Dartmouth College and NIH funding from P20-GM113132 and 5R01NS112365. Use of the spinning disk confocal microscope was made possible by the following funding sources: NIH award S10OD032310 to Dr. Yashi Ahmed, the Light Sciences Light Microscopy Facility at Dartmouth, BioMT through NIH grant P20-GM113132, DartCF through NIH grant P30-DK117469, as well as a scholarly innovation award from Dartmouth College, and the Dartmouth Cancer Center with NCI Cancer Center Support Grant P30 CA023108.

## 5. AUTHOR CONTRIBUTIONS

Conceptualization: A.S.A and M.B.H. Methodology: A.S.A and M.B.H. Investigation: A.S.A,

M.W.M. Writing (original draft): A.S.A. and M.B.H. Writing (reviewing and editing): A.S.A, M.W.M, and M.B.H. Funding acquisition: A.S.A. and M.B.H. Project administration and supervision: M.B.H.

## 7. SUPPLEMENTAL INFORMATION

- Document S1. Figures S1–S6.

## 8. STAR★ Methods

### 8.1. KEY RESOURCES TABLE

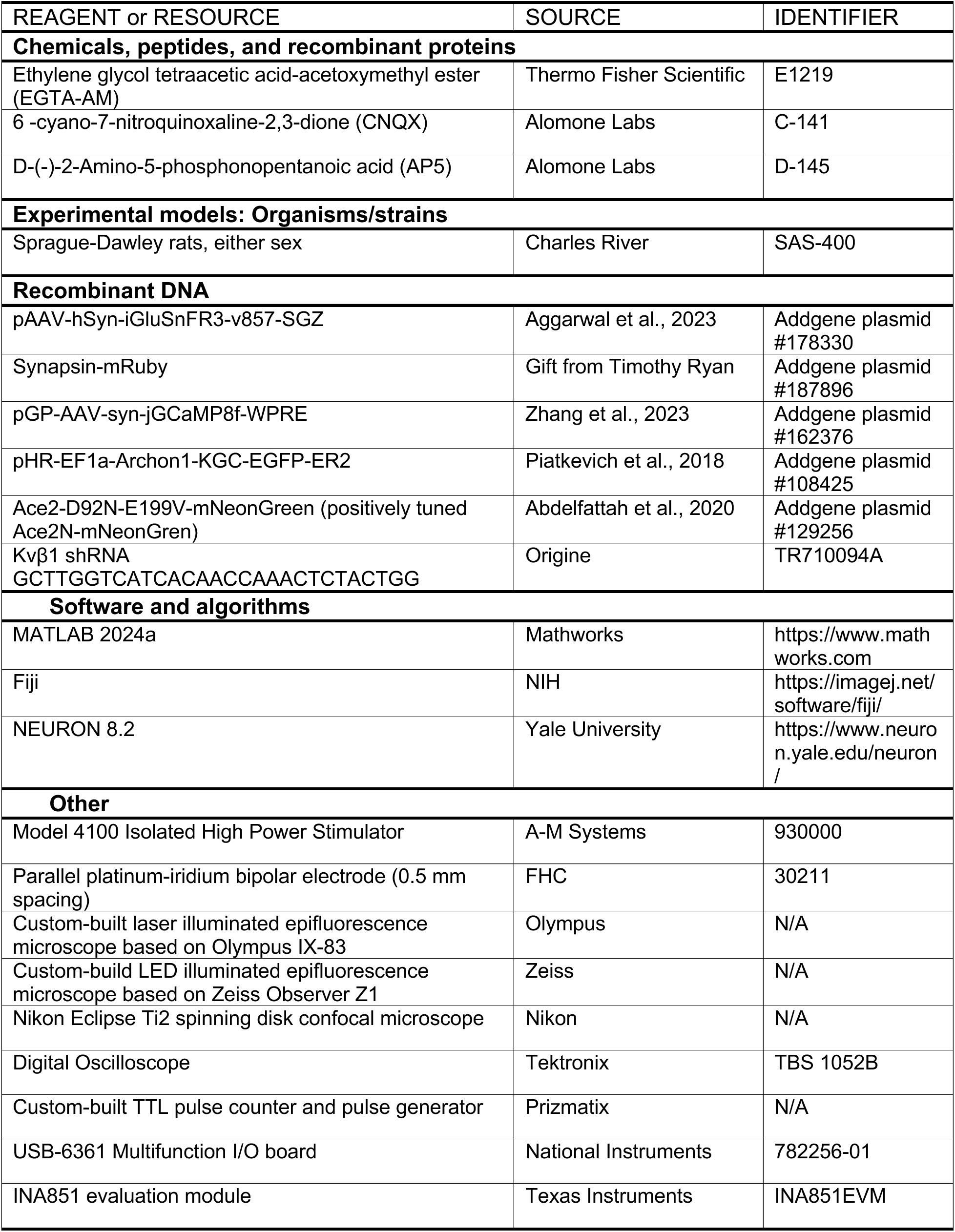

### 8.2. RESOURCE AVAILABILITY

#### Lead contact

Further information and requests for resources and reagents should be directed to Michael Hoppa

#### Materials availability

This study did not generate new, unique reagents.

#### Data and code availability

All data and analysis code reported in this paper will be shared by the lead contact upon request.

### 8.3. EXPERIMENTAL MODEL AND SUBJECT DETAILS

Experiments were conducted in primary hippocampal neurons cultured from postnatal day 0 Sprague-Dawley rats of either sex, using the protocol described below.

### 8.4. METHOD DETAILS

#### Cell culture and transfection

Hippocampal CA1–CA3 regions were dissected, excluding the dentate gyrus, and the tissue was dissociated with bovine pancreas trypsin (5 min at room temperature) and plated on polyornithine-coated glass coverslips within 6 mm diameter cloning cylinders, as described previously ^66^. Cultures were incubated at 37° C and 95% air/5% CO_2_ humidified incubator On DIV 5 or 6, cultures were sparsely transfected with plasmid DNA using Ca^2+^ phosphate-mediated gene transfer and a total of 6–7 µg DNA per culture. All genetically encoded fluorescent indicators, along with other reagents, can be found in the Key Resources Table. All experiments were conducted on neurons between DIV 14 to 20 and repeated on neurons from three independent cultures. The protocols used were approved by the Institutional Animal Care and Use Committee at Dartmouth College and conform to the NIH Guidelines for the Care and Use of Animals.

#### Live cell imaging and stimulation

Neurons were imaged on custom-built inverted epifluorence microscopes with a 3D printed rapid-switching laminar flow perfusion and stimulation chamber placed over the glass coverslip (**Figure S1**). The 3D printed chamber held a volume of 80 µL with a 7 x 0.5 mm (cross-section) perfusion channel spanning the 18 mm chamber width and two pairs of parallel, 5 mm long platinum-iridium (Pt-Ir) electrodes placed 6.5 mm apart. Uniform E-fields were generated by applying isolated current pulses (A-M Systems Model 4100) between either of the two pairs of electrodes, which were positioned such that the 6 mm diameter neuron cultures were centered between them.

During imaging, neurons were perfused at 300–400 µL/min in a modified Tyrode’s solution consisting of (in mM): 119 NaCl, 2.5 KCl, 1.3 CaCl_2_, 2.7 MgCl_2_, 25 HEPES, 30 glucose (Sigma-Aldrich). For experiments with altered extracellular K^+^ concentration, we used Tyrode’s with modified KCl and equimolar changes in NaCl to maintain osmolarity. In a subset of experiments, we used Tyrode’s with 3.5 mM extracellular Ca^2+^. To silence network activity and indirect responses to stimulation, we added 10 µM 6-cyano-7-nitroquinoxaline-2,3-dione (CNQX) and 50 µM D-(−)-2-Amino-5-phosphonopentanoic acid (AP5), both from Alomone Labs. Temperature was held at ∼35°C using a custom-built objective heater.

High speed imaging of glutamate, calcium, and voltage indicators was performed on an Olympus IX-83 microscope with a 40X/1.35 numerical aperture (NA) oil immersion objective (UApo-N-40X-O340-2). Images were captured with an Andor IXON Ultra 897 electron-multiplying charge coupled device (EM-CCD) camera (Oxford Instruments Andor), with imaging and stimulation protocols described below for each indicator. The EM-CCD camera was liquid cooled (EXOS) to −80°C. Stimulation timing was synchronized with camera frame acquisition timings using either a custom-built TTL pulse counter and generator (Prizmatix Ltd.) or using an NI USB 6361 multifunction I/O board, controlled in MATLAB (R2024a, The Mathworks, Inc.) using Wavesurfer (v1.0.6, Janelia). Excitation light was provided via OBIS lasers (Coherent) with the following wavelengths and filter sets (Chroma): 488-nm laser with ET470/40×, ET525/50m, and T495lpxr filters (iGluSnFR3/jGCaMP8f); 561-nm laser with ET560/40×, ET630/75m (synapsin-mRuby), and T585lpxr filters; and 637-nm laser with ZET635/20×, ET655lpm, and ZT640rdc filters (Archon1).

##### Subthreshold modulation of glutamate release and calcium influx (iGluSnFR3/jGCaMP8f)

For iGluSnFR3 and jGCaMP8f, images were captured with a sampling rate of 100 Hz (9.7 ms exposure time) and 2–5 mW power (488-nm laser). The glutamate sensitive domain of iGluSnFR3 is on the extracellular side of the plasma membrane, so fluorescence from the iGluSnFR3-expressing neuron is potentially sensitive to glutamate spillover from APs in nearby untransfected neurons activated by a spatially uniform E-field stimulus. Therefore, for iGluSnFR3 imaging, we used a more focal stimulus to evoke action potentials (APs) in the imaged neuron using parallel Pt-Ir bipolar electrodes (0.5 mm spacing; #30211, FHC, Inc.) positioned near its cell body. Rectangular, 1 ms monophasic pulses were applied with the amplitude adjusted to approximately 20% above threshold, which varied between cells and depending on the precise electrode position (1–4 mA). Cytosolic calcium measurements do not suffer from artifacts related to activation of non-transfected neurons, so for jGCaMP8f imaging, we instead used uniform E-field stimulation on the perpendicular set of electrodes to evoke APs.

We measured the effect of subthreshold, uniform E-field on AP-triggered glutamate release or cytosolic calcium influx by evoking a 2 Hz train of 60 APs and applying a 10 sec long uniform E-field after the first 20 APs, starting 250 ms before the 21^st^ AP and ending 250 ms after the 40^th^ AP. This allowed for within-trial comparison of glutamate release or calcium influx before, during, and after subthreshold E-field stimulation for each synapse. Recordings at each subthreshold E-field amplitude were repeated 2–3 times with at least 1 minute between trials. For iGluSnFR3 and jGCaMP8f measurements, neurons were cotransfected with synapsin-mRuby to identify presynaptic boutons for subsequent analysis. The synapsin-mRuby signal was imaged at the start and end of experiments.

For the EGTA experiments with iGlusnFR3 (**Figure 4**), we loaded neurons with EGTA by incubating in Tyrode’s buffer with 100 µM EGTA-AM (Thermo Fisher Scientific) for 2 min and then perfusing for 10 min in control Tyrode’s. EGTA-AM stock aliquots were prepared at 20 mM in dimethylsulfoxide (DMSO).

We also measured how facilitation by subthreshold uniform E-fields interacted with paired pulse facilitation of glutamate release with iGluSnFR3 (**Figure 4**). For these experiments, a single trial consisted of evoking a single AP, followed by a 5 sec delay and two APs at 20 Hz. This protocol allowed separate analysis of peak iGluSnFR3 response to both a single and paired pulse, although we found the high SNR and fast kinetics of iGluSnFR3 imaged at 100 Hz allowed separation of the peaks for the 20 Hz paired pulse stimulus used. We alternated between control trials (single and paired pulse stimuli with no subthreshold E-field) and E-field trials in which a 2.1 sec duration uniform E-field was applied starting 2 sec before both the single and paired pulse stimuli. For these experiments, the E-field amplitude was measured immediately before imaging and the current intensity was adjusted to generate a 50 V/m E-field, which required 1–1.6 mA depending on exact perfusion level. Because axonal orientation varies between neurons in each culture, whether a given E-field direction was depolarizing or hyperpolarizing for the axon in the imaged field of view (FOV) was effectively random. Thus, we identified a facilitating E-field direction by first applying a 10 sec subthreshold E-field during a 2 Hz AP train (protocol described earlier) with both polarities (+50 and −50 V/m). This E-field direction was then used in the paired pulse trials.

##### Axonal AP waveform and polarization measurements (Archon1)

To measure the effect of subthreshold uniform E-fields on axonal membrane potential and AP waveform, we imaged Archon1 in individual axonal branches with a sampling rate of 2 kHz (0.483 ms exposure time) and 50–70 mW laser power (637-nm laser). To achieve this frame rate, the camera was run in cropped sensor mode (10 MHz readout, 0.5 ms pixel shift speed in a 10 × 36 µm FOV) with an intermediate image plane mask (Optomask, Cairn Research) to block light on nonrelevant pixels, as described previously ^34^. For AP waveform measurements, we imaged trains of 40 APs evoked at 2 Hz. In subthreshold E-field trials, we applied a 10 sec long uniform E-field (±1 mA, ∼55 V/m) during the latter 20 APs (starting 250 ms before the 21^st^ AP). Here, we evoked APs using 1 ms, 15 mA (∼825 V/m) current pulses applied to the perpendicular set of E-field electrodes, since the membrane potential/voltage-sensitive fluorescence is not affected by activation of non-transfected background cells. We then compared the averaged AP waveforms of the first 20 APs and the last 20 APs (during E-field), providing within-trial controls. In the same cells, we measured the membrane polarization generated by the subthreshold E-field alone using 10, 500 ms E-field pulses applied at 1 Hz with multiple current intensities between −2 and 2 mA. Calibrations of polarization to AP amplitude used the control AP waveform measurement from the last trial collected.

##### Wide FOV polarization imaging of pAce2N-mNeon

Finally, to image steady-state polarization across the morphology of a single neuron (**Figure 1**), we instead used a Zeiss Observer Z1 microscope optimized for high spatial resolution imaging over a large field of view using more uniform LED illumination and a complementary metal-oxide semiconductor (CMOS) camera. To image pAce2N-mNeon fluorescence, we used an EC Plan-Neofluar 40x 1.3 NA oil immersion objective. For illumination, we used a 475/28 nm diode and the SPECTRA solid-state light engine (Lumencor), and we captured images on an ORCA-Fusion Digital CMOS Camera (Hamamatsu). Perfusion and temperature control were the same as in the experiments described above. Polarization responses were measured by imaging at 20 Hz over a 374 × 374 µm FOV using the full chip (2304×2304 pixels, with 0.1625 µm pixels and 2×2 binning). Each trial consisted of twenty, 500 ms long E-field pulses applied at 2 mA (∼110 V/m). This was repeated with 4 E-field directions (0°, 90°, 180°, and 270°) by altering the polarity and pair of electrodes (horizontal or vertical). The stimulus-aligned movies were then averaged for subsequent processing and analysis. Immediately following the functional imaging experiment, structural imaging was conducted on the same neuron using a Nikon Eclipse Ti2 spinning disk confocal microscope with a 60x, 1.42 NA oil immersion objective (Plan Apo Lambda D). A tiled z-stack (0.1625 µm lateral resolution, 0.3 µm z-step) was acquired encompassing the full neuronal morphology.

### Morphological reconstruction and neuron modeling

We used the high-resolution tiled z-stack of the pAce2N-mNeon-expressing excitatory neuron imaged during subthreshold uniform E-field stimulation to build a morphologically realistic neuron model. This model allowed us to replicate the experiment *in silico* in the NEURON simulation software (v8.2) ^67^. Morphology tracing of the entire dendritic and axonal arbor (shown in **Figure 1B**) was conducted in ImageJ using the simple neurite tracer (SNT) plugin ^68^.

Simulations were controlled in MATLAB (R2024a, The Mathworks, Inc.) using our previously developed framework for defining, running, and importing data from NEURON simulations of extracellular E-field stimulation of single neurons^22,69^. After importing the morphology, we distributed sets of ion channel conductances in the soma, dendrites, and axonal compartments, with the same models and channel densities previously used for the model of layer 5 thick-tufted neocortical pyramidal neurons (cADpyr232) from P14 rats by the Blue Brain Project^22,70^. We assigned the conductances of the pyramidal neuron basal dendrites to all dendritic compartments of the modeled cultured hippocampal neuron, which lacks clear apical and basal dendrites. Uniform E-field stimulation was simulated by assigning extracellular potentials to each compartment using the `extracellular` mechanism and scaling them uniformly in time based on the temporal waveform. Replicating the experiment, we simulated 500 ms long E-field (100 V/m) in each direction. Simulations were run with a 25 µs time step and the backward Euler solver. The model was spatially discretized with isopotential compartments no longer than 2.5 µm, giving 10,318 compartments. Simulations were run on a Macbook Pro (10-core M1 Pro, 2021) with 32 GB RAM.

### E-field calibration

We used a custom-built E-field recording probe using a pair of Ag/AgCl or PtIr wires (0.5 mm diameter) mounted on a 3D printed holder with interelectrode distance of 4 mm. Recorded voltage difference was differentially amplified (gain of 13x) using INA851 instrumentation amplifier with its accompanying evaluation board and recorded using a National Instruments USB-6361 multifunction I/O board, controlled in MATLAB using Wavesurfer (v1.0.6, Janelia). The E-field amplitude was calculated by dividing the measured voltage difference (pre-amplification) by the interelectrode distance. For calibration before imaging experiments, we applied train of five, 100 ms rectangular pulses at three intensities (0.1, 0.2, and 0.3 mA). With the stimulus-averaged pulse at each intensity, we then extracted the steady state value (mean of final 50 ms) at each intensity and fit the steady state E-field vs. applied current to a line with least squares regression. The slope (in V/m per mA) was used to set the current intensity achieving the desired E-field for subsequent stimulation trials. Variations in perfusion and bath height between sessions introduced variability in current density, and therefore E-field magnitude, which could be mitigated using a glass cover in experiments not requiring additional electrodes to be positioned near the imaged neuron (**Figure S2**). Example recordings are shown in **Figure S2B**. We verified the E-field was spatially uniform by moving the probe laterally and in depth within the bath and found negligible change in E-field amplitude (data not shown).

We used unmyelinated cultured neurons from rats, which allowed for ideal optical access for high-resolution imaging of signals from subcellular compartments. However, these neurons likely have reduced polarization sensitivity (i.e., mV polarization per V/m) relative to mature, human neurons due to their smaller diameters and lack of myelination ^22^. Therefore, we applied a range of E-field intensities (0.1 – 1 mA; 5.5 – 55 V/m) that generated axonal polarization, based on our AP-calibrated optical polarization measurements, encompassing the range expected for tDCS. Axonal polarization in our cultured neurons was measured using our AP-calibrated optical polarization measurements, and expected axonal polarization for tDCS was based on our previously published biophysical modeling (0.3–5 mV) ^23^. It is worth nothing the subthreshold stimulation never evoked APs at the E-field intensities used. Also, it is unlikely the inhibiting E-field directions blocked AP propagation, which we previously showed occurs with virtually no propagation failures at baseline ^71^, based on the consistent jGCaMP8f AP-evoked responses in all boutons.

### 8.5. QUANTIFICATION AND STATISTICAL ANALYSES

#### Image analysis and statistics

All image and statistical analyses were done using custom software written in MATLAB. To extract fluorescence time series, regions of interest (ROIs) were created in ImageJ (NIH) and loaded into MATLAB for downstream processing. Values in text and figures are mean ± standard error (SEM) unless otherwise stated. When data are shown as averaged traces with error bars, error is SEM unless otherwise stated. For paired comparisons, statistical significance was assessed using a two-sided Wilcoxon signed rank test for non-normal distributions or two-sided paired t-test for normally distributed data. Normality was assessed using the Kolmogorov-Smirnov test. Below, we describe the additional image processing and statistical analyses for specific experiments.

##### Subthreshold modulation of glutamate release and calcium influx (iGluSnFR3/jGCaMP8f)

To extract fluorescence signals within single presynaptic boutons, 2.4 µm diameter circular ROIs were manually selected using the synapsin-mRuby signal. For analysis of peak AP-triggered responses, we computed baselined-normalized change in fluorescence (Δ*F*/*F*_0_) responses using the 15 ms preceding each AP for the local baseline (*F*_0_). Peaks were extracted from the 50 ms after each stimulus. Boutons on branch points or overlapping with another axonal branch or neighboring boutons were excluded. We included data from cells with at least 10 in-focus boutons for the duration of experiment that all pass the following inclusion criteria related to fluorescence signal and physiological stability: 1) Less than 15% change in baseline from first trial to final trial; 2) No trial with stimulus-averaged peak response of the 20 control APs (before subthreshold E-field stimulation) that deviates by more than 60% from the average across trials, to exclude synapses with rundown or instability due to imaging conditions. We identified presynaptically “silent” synapses as synapses with evoked responses for all APs below a threshold of 1 standard deviation above pre-stimulus baseline fluorescence (**Figure 2F**).

Since cultured neurons do not have morphologies with consistent branch orientations, the relationship between E-field direction and polarity of voltage changes was essentially random from cell to cell. For this reason, +1 mA could be facilitating in the imaged boutons of one neuron and inhibiting in another, and this polarity dependence is essentially arbitrary and unknown without simultaneous voltage recordings. Therefore, to analyze modulation of glutamate release or calcium influx across cells, we grouped data at each intensity by determining the direction producing strongest modulation (facilitating or inhibiting) for each bouton at the highest intensity tested, defined as the largest absolute percent change in mean peak iGluSnFR3 or jGCaMP8f response during relative to before application of the subthreshold E-field. If this relative change was positive (facilitating), this direction was assigned the positive polarity (e.g., +1 mA), and if this relative change was negative (inhibiting), this direction was assigned the negative polarity (e.g., −1 mA).

After this polarity sorting procedure, we tested the effect of subthreshold E-field intensity and trial phase (before, during, and after 10 sec E-field) on peak responses using two-way repeated measures ANOVAs with post-hoc Tukey-Kramer tests for multiple comparisons. The polarity sorting procure necessary for population level analysis could cause random variations in peak amplitudes to appear as modulation during the E-field, so we also conducted within bouton analysis to identify responding synapses using all the peak responses without averaging. For this analysis, we used Wilcoxon ranked sum tests to compare the distribution of peak response amplitudes before vs. during (or after) the 10 sec E-field in each bouton, which consisted of 40 AP-triggered responses per trial phase for each E-field intensity (2 trials per intensity). To correct for multiple comparisons, we used the Benjamini and Hochberg procedure for controlling for the false discovery rate of a family of hypothesis tests^72^ with a conservative false discovery rate of 1%.

##### Axonal AP waveform and polarization measurements (Archon1)

For Archon1 recordings from individual axonal branches, we extracted fluorescence signals within manually drawn ROIs encompassing the imaged axonal branch. Stimulus-triggered action potential waveforms were averaged and relative change in fluorescence (Δ*F*/*F*_0_) was computed using a baseline window of 15 ms preceding each stimulus. AP amplitude and full width half max (FWHM) was extracted from stimulus-averaged axonal AP waveforms that were upsampled from 2 kHz to 10 kHz using cubic spline interpolation. For measurements of polarization with 500 ms subthreshold E-fields, mean Δ*F*/*F*_0_ responses were computed using a baseline window was 200 ms and high-frequency noise was filtered out using a quadratic Savitsky-golay filter with a 20 ms window. Polarization time constants (*τ*_*r*_) were calculated using nonlinear least square fitting of the 20 ms immediately after the start of the subthreshold E-field to the equation *y* = *Ae*^−*t*/*τ*^ (**Figure S4**). Parameter fits with *R*^2^ < 0.7 and SNR < 4 were excluded from further analysis.

##### Wide FOV polarization imaging (pAce2N-mNeon)

The data from this experiment consisted of functional recordings (20 Hz) of membrane polarization in response to 500 ms E-field pulses and high-resolution 3D structural imaging of the same neuron. To generate the maps of steady-state polarization in **Figure 1C** and **Figure S3** we used the following image processing steps, parts of which were adapted from steps used for mapping subcellular AP propagation in high-speed voltage imaging movies ^73,74^. First, a stimulus-averaged movie was computed for the 20 E-field pulses in each recording. Then, movies were denoised using a spatial gaussian filter (width of 2 pixels, 9×9 filter) followed by a principal components analysis (PCA)-based temporal filtering step in which the top 4 PCA eigenvectors were retained within each pixel. Mean steady state change in fluorescence was then computed using the mean change in fluorescence (Δ*F*/*F*_0_) of the last 400 ms of the 500 ms E-field pulse, using a 450 ms baseline. This steady-state polarization image was upsampled and coregistered with the max z-projection of the high-resolution 3D structural image using the `imregtform` MATLAB function with an affine transformation. The lower-resolution polarization data were then mapped onto the higher-resolution structural image by masking the steady-state polarization image with a user-defined threshold pixel intensity from the structural image. Pixels with maximum SNR below 3 across all trials (E-field directions) were also masked to minimize conflation of weak polarization with low signal. Finally, the masked polarization images were visualized using the `surf` function in MATLAB.

## 10. Supplementary Figures

**Figure S1, related to Figure 1.**
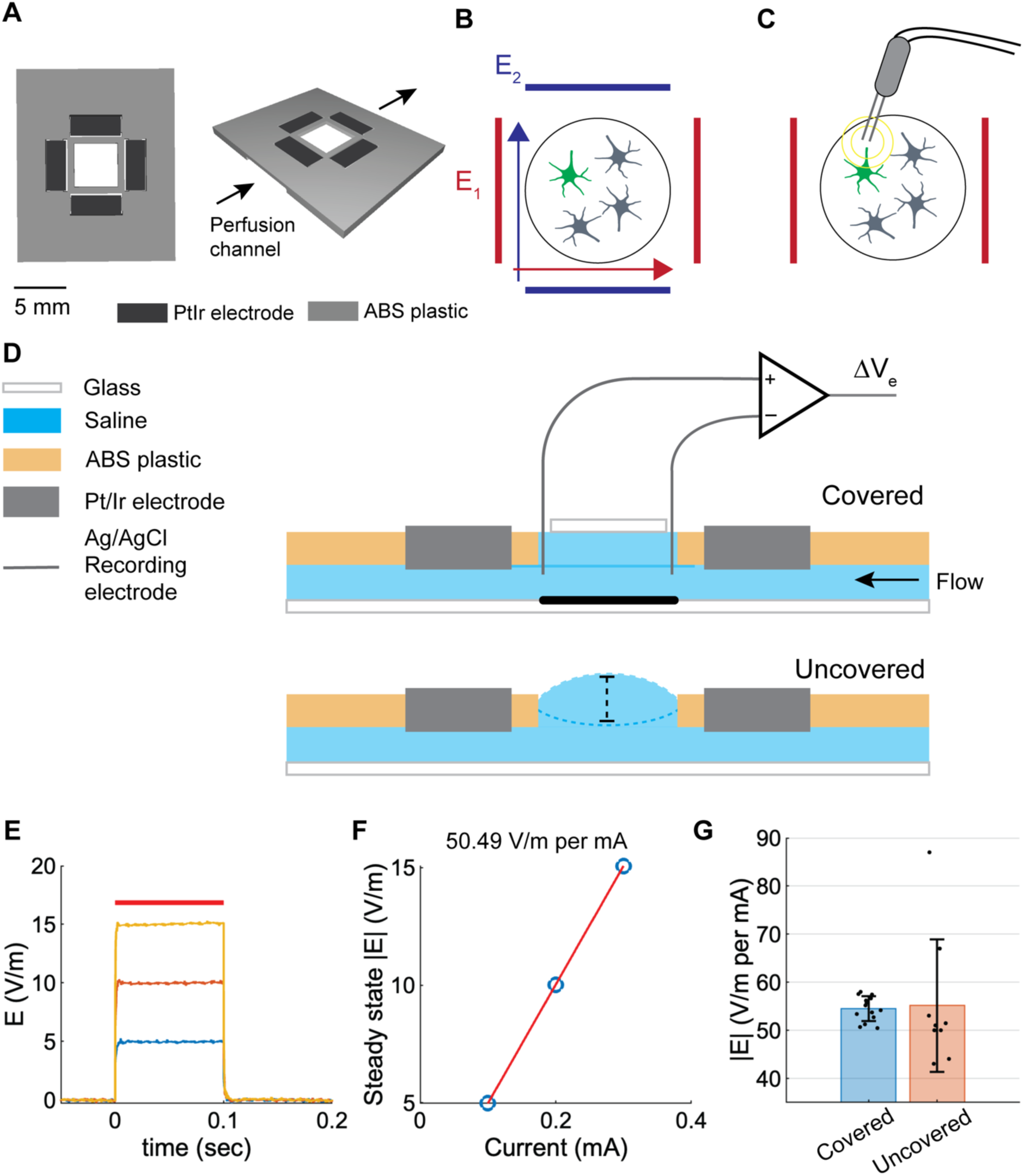
Stimulation configurations and electric field recording setup and effect of bath height. **A)** CAD model of 3D printed live cell imaging chamber for E-field stimulation and perfusion. Electrodes were exposed on underside of chamber, above the perfusion channel to pass current across the neuron cultures. Neurons were cultured in 6 mm cylinders on glass coverslips, which were positioned beneath the central window of the chamber. **B)** Cartoon version of stimulation chamber showing uniform E-field stimulation with arbitrary orientation using both pairs of parallel PtIr electrodes. **C)** Cartoon showing second stimulation configuration for focal stimulation of imaged neuron with bipolar PtIr electrode during application of uniform E-field with one pair of parallel PtIr electrodes. **D)** Schematic of two configurations for imaging experiments: covered perfusion, which provides more stable bath height while still allowing for E-field recording electrodes (top) and uncovered perfusion, necessary for lowering in and freely positioning bipolar stimulation electrodes to more focally activate individual neurons. **E)** Example E-field recordings using E-field calibration protocol before imaging experiments. E-field pulses with 100 ms duration were applied in trains consisting of 5 pulses at 5 Hz, baseline corrected, and averaged for each intensity (0.1, 0.2, and 0.3 mA). **F)** Steady state E-field amplitude at each intensity shown in B) with linear fit overlaid (red). **G)** E-field calibration measurements showing variability in either configuration with perfusion. Measured E-field was 54.5 ± 2.6 V/m per mA (*n* = 14, mean ± STD) for covered and 55.2 ± 13.8 V/m per mA (*n* = 9) for uncovered stimulation chamber. Individual data points are from measurements using the same electrode chamber in separate sessions.

**Figure S2, related to Figure 1.**
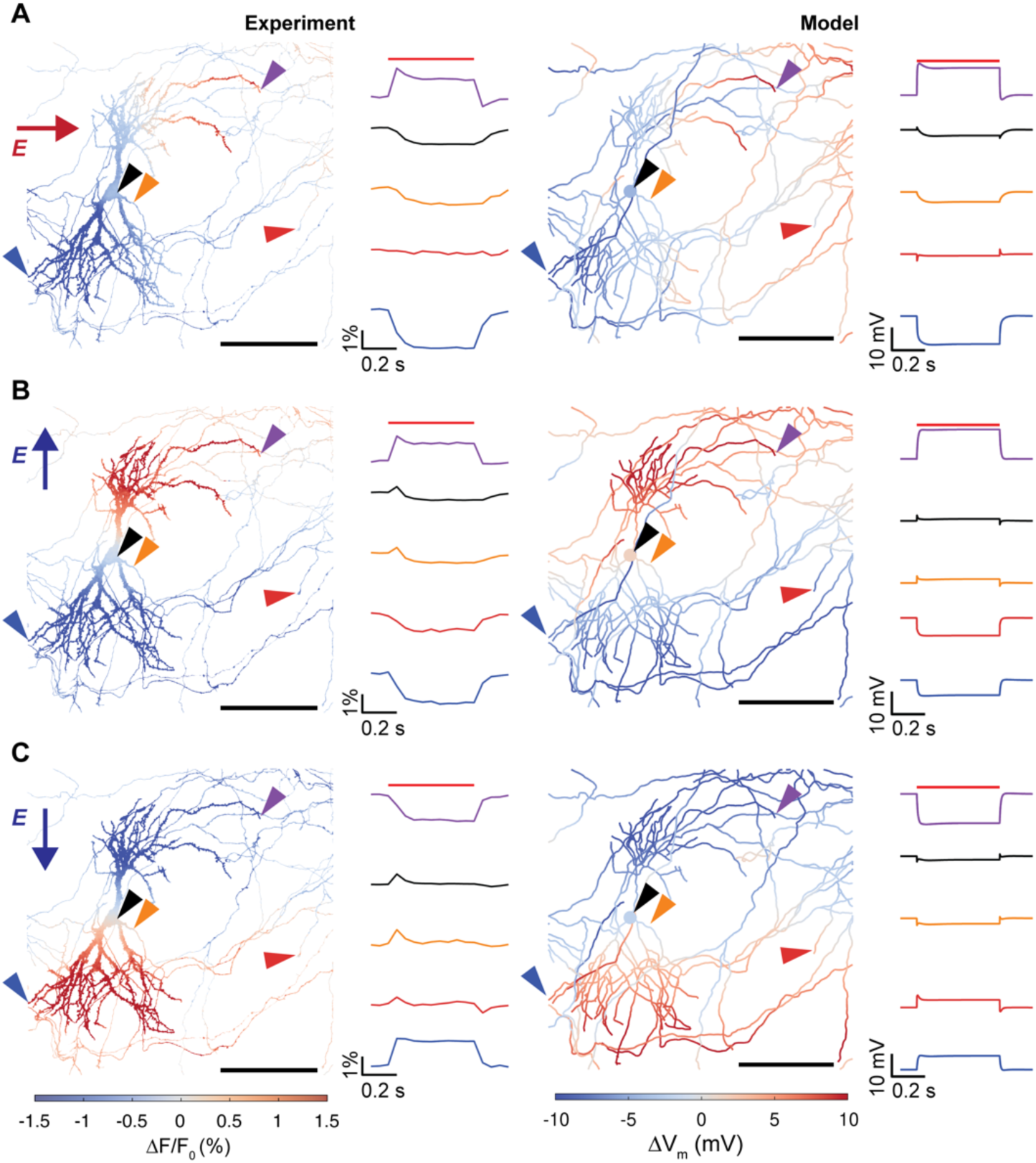
Effect of E-field direction on subcellular polarization distribution. Measured (left side) and simulated (right side) membrane polarization for 500 ms subthreshold E-field (2 mA, ∼110 V/m) directed right (A), up (B), and down (C). As in Figure 1, for optical measurements of pAce2N-mNeon (left side), colors indicate steady-state change in fluorescence and fluorescence traces from example ROIs (right inset). For simulation (right side), colors indicate simulated steady-state change in membrane voltage and voltage traces from same ROIs as in experiment (right inset).

**Figure S3, related to Figure 1.**
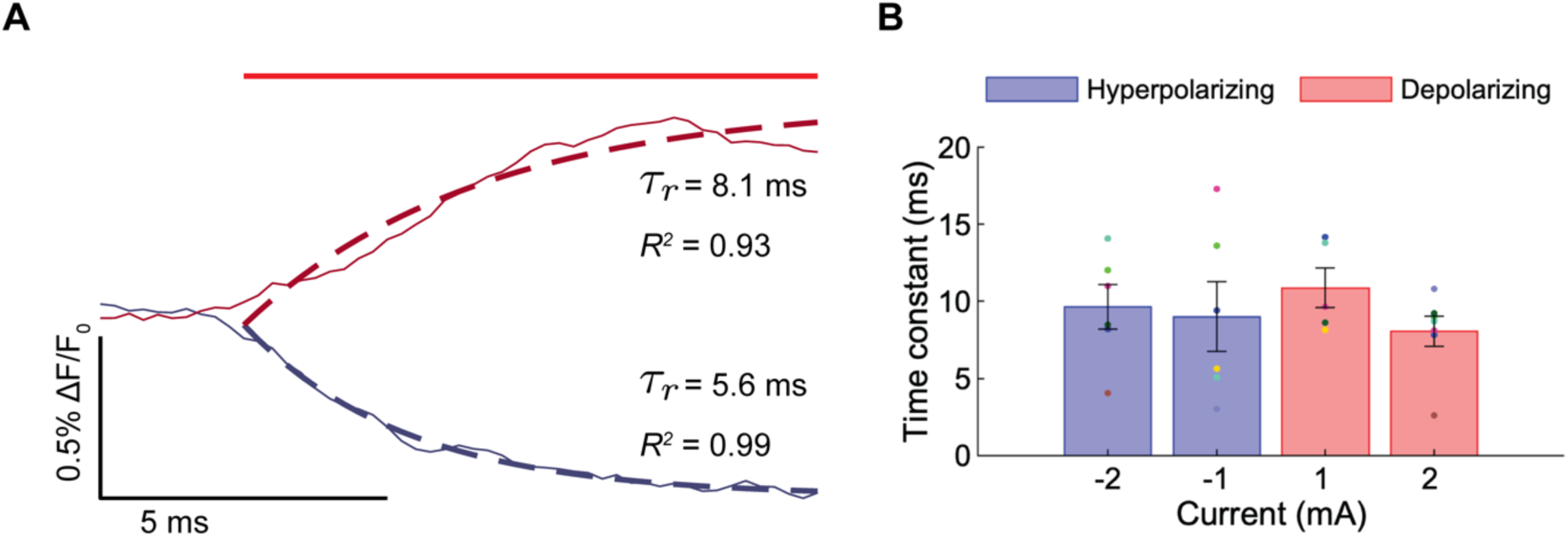
Axonal membrane polarization time constant measured with Archon1. **A)** Example fluorescence traces (solid lines) showing initial response (20 ms) to depolarizing (red) and hyperpolarizing (blue) uniform E-field (500 ms total duration), with overlaid monoexponential fits (dashed lines). Fluorescence traces filtered with Same axon as in Figure 1. **B)** Estimated polarization time constants measured at each current intensity. Excluded parameter fits with *R*^2^ < 0.7 and SNR < 4. Time constants were not significantly different between stimulation intensities (one-way ANOVA); combined across intensities time constants were 9.2 ± 1.1 ms (*n* = 8, mean ± SEM).

**Figure S4, related to Figure 1.**
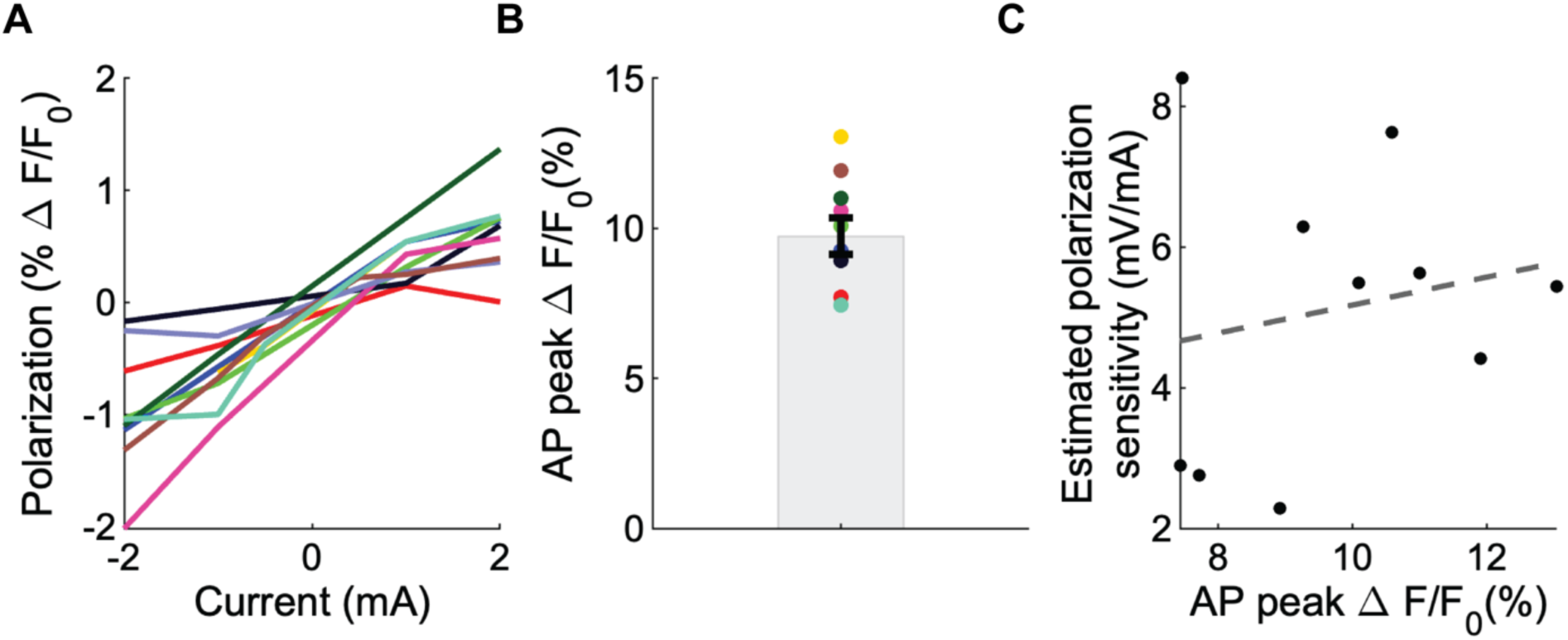
Calibrating axonal E-field polarization to AP amplitude. **A)** Steady-state polarization (as fluorescence change) within one axonal region of interest per neuron plotted against applied current intensity**. B)** Mean AP amplitude was 9.7 ± 0.6% Δ*F*/*F*_0_ (mean ± SEM, n = 10 cells). **C)** Estimated polarization sensitivity (assuming an axonal AP amplitude of 120 mV) plotted against AP peak for each axon, with linear fit overlaid (*R*^2^ = 0.035, *p* = 0.6).

**Figure S5, related to Figure 4.**
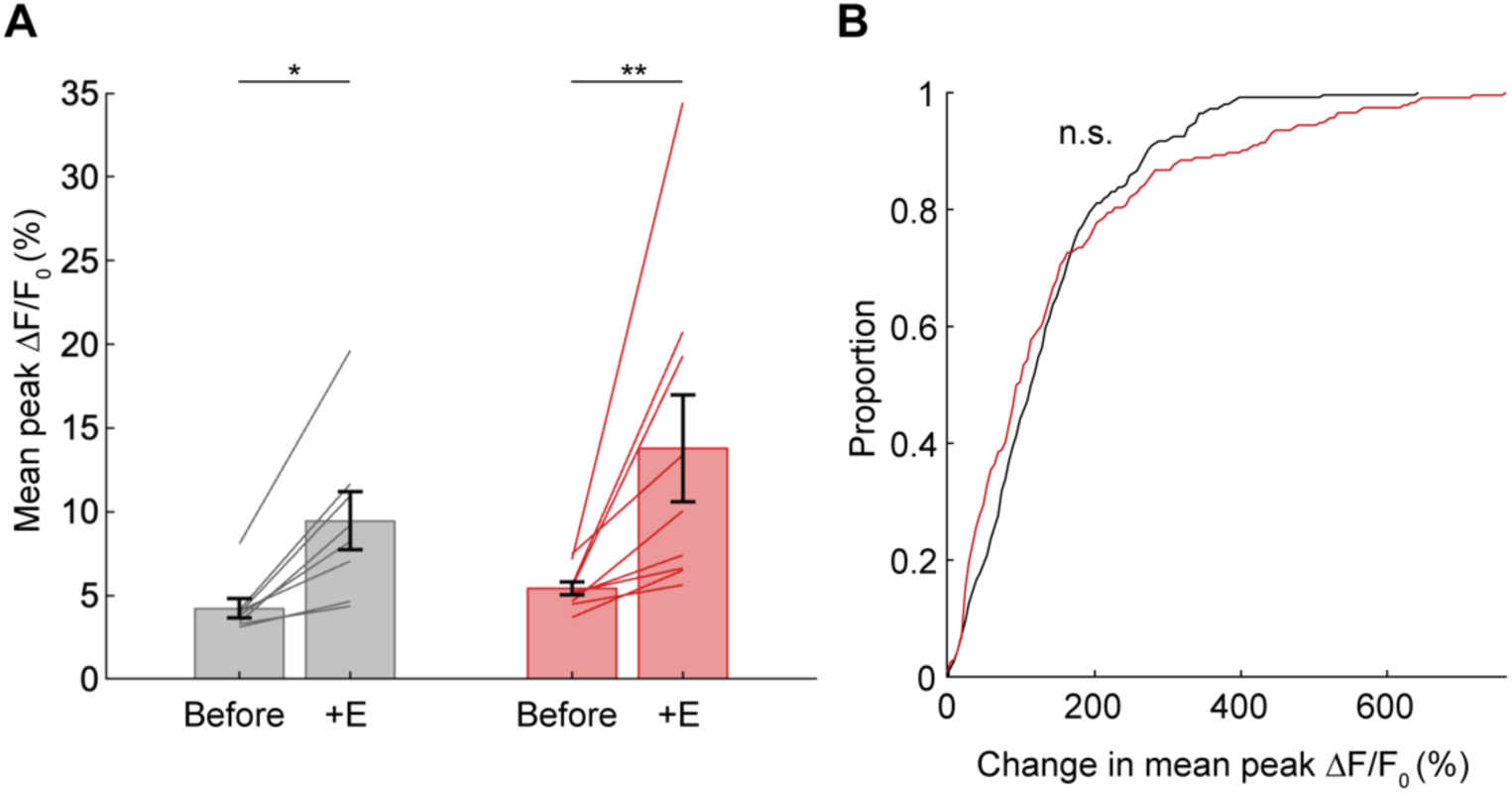
Knockdown of KvBeta1 has no effect on facilitation of glutamate release by subthreshold E-field. **A)** Mean peak evoked iGluSnFR3 Δ*F*/*F*_0_ before and during 1 mA, 10 sec DC E-field in wild-type (WT) and K_v_β1 shRNA transfected neurons (n = 254 synapses, 8 cells for WT; n = 234 synapses, 10 cells for KD). Main effect of genotype (p = 0.22) and genotype x stimulation interaction not significant (p = 0.36) for two-way RM ANOVA (* p < 0.05, ** p < 0.01 post-hoc Tukey-Kramer for effect of E-field within genotype). **B)** Cumulative distribution plot of percent change in mean of peak evoked glutamate release (Δ*F*/*F*_0_) during 1 mA, 10 sec DC E-field relative to before stimulation across all boutons of WT and K_v_β1 shRNA transfected neurons.

**Figure S6, related to Figure 4.**
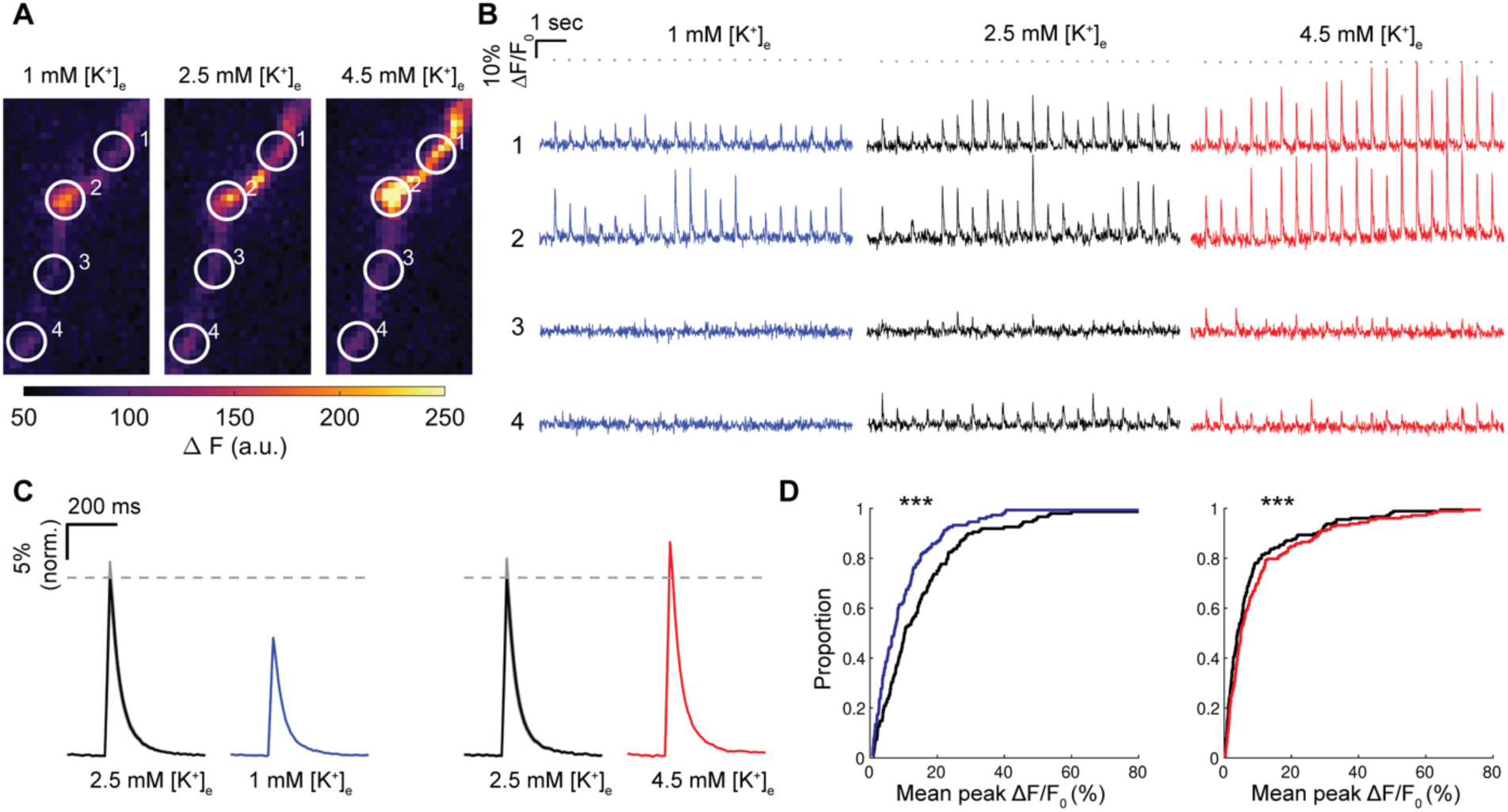
Depolarizing rest potential facilitates, and hyperpolarizing inhibits, evoked glutamate release. **A)** Mean AP-evoked change in fluorescence in example axon with four boutons (based on synapsin-mRuby signal) in 1 mM, 2.5 mM, and 4.5 mM [K^+^]_e_. Tyrode’s buffer. **B)** Fluorescence traces from ROIs in A) during 2 Hz AP train in each [K^+^]_e_ condition. **C)** Stimulus-averaged responses (mean ± SEM) across all cells in 2.5 mM (control) and either 1 mM (*n* = 148 boutons, 5 cells) or 4.5 mM [K^+^]_e_ (*n* = 123 boutons, 5 cells), normalized to mean in 2.5 mM [K^+^]_e_. **D)** Cumulative distribution plots of peak evoked responses (Δ*F*/*F*_0_) in 2.5 mM (control) and either 1 mM (left) or 4.5 mM [K^+^]_e_ (right), *** *p* < 0.001, Wilcoxon signed-rank.

